# Using BONCAT To Dissect The Proteome Of *S. aureus* Persisters

**DOI:** 10.1101/2024.11.01.621614

**Authors:** Eva D. C. George Matlalcuatzi, Thomas Bakkum, Pooja S. Thomas, Stephan M. Hacker, Bogdan I. Florea, Bastienne Vriesendorp, Daniel E. Rozen, Sander I. van Kasteren

**Author notes:** To whom correspondence should be addressed: DER; SIvK.

## Abstract

Bacterial persisters are a subpopulation of cells that exhibit a transient non-susceptible phenotype in the presence of bactericidal antibiotic concentrations. This phenotype can lead to the survival and regrowth of bacteria after treatment, resulting in relapse of infections. As such, it is also a contributing factor to antibacterial resistance. Multiple processes are believed to cause persister formation, yet identifying the proteins expressed during the induction of the persister state has been difficult, because the persister-state is rare, transient and does not lead to genetic changes. In this study, we used Bio-Orthogonal Non-Canonical Amino Acid Tagging (BONCAT) to label, and retrieve, the proteome expressed during the persister state for different strains of methicillin-resistant *Staphylococcus aureus*. After incubating antibiotic-exposed bacteria with the methionine ortholog L-azidohomoalanine to label the proteins of persister cells, we retrieved labeled proteins using click chemistry-pulldown methodology. Analysis of the retrieved proteome fraction of Methicillin resistant *Staphylococcus aureus* (MRSA) and Vancomycin resistant *Staphylococcus aureus* (VRSA) under challenge with β-lactam and fluoroquinolone antibiotics with Label Free Quantification - Liquid chromatography mass spectrometry (LFQ-LCMS) based proteomics reveals the upregulation of proteins involved in stringent response, cell wall biosynthesis, purine metabolism, ppGpp biosynthesis, two component systems (TCS), lipid metabolism, ABC transporters, D-alanine biosynthesis and L-proline degradation. Conversely, we observed a decline of proteins associated with amino acid biosynthesis and degradation, protein biosynthesis, protein modification, and carbohydrate metabolism, among others. These findings indicate that modification of translational activity in persister cells enables bacterial cells to induce an active defense to survive antibiotic pressure.

## Introduction

Persisters are a subpopulation of bacteria that adopt a dormant slow/no-growth-state after exposure to lethal antibiotic concentrations.[1, 2] These cells are able to survive during antibiotic challenge and then resume growth after antibiotic removal, causing relapse. They are also more likely to evolve antibiotic resistance.[3] Persisters are therefore a key obstacle to bacterial clearance during treatment of infections.[4] The persister phenotype is apparently ubiquitous in microbes and has been detected in fungi[5], Gram-positive[6], and Gram-negative species[7], including major pathogens such as *Mycobacterium tuberculosis* [8, 9], *Escherichia coli* [10, 11], *Pseudomonas aeruginosa* [12, 13], *Salmonella typhimurium*[14], and *Staphylococcus aureus*.*[15, 16]* However, the mechanisms underlying this phenotype remain uncertain.

Research on how bacteria enter, maintain, and depart the persister state has been challenging.[2] One of the major reasons for this is that the fraction of persister cells in a treated population is very small, comprising only ∼0.001-0.1% of cells. [17, 18] Identifying persistence mechanisms is therefore difficult because it requires enrichment and isolation of this rare minority. [19] A second complicating factor is that the persister phenotype is transient, with the bacteria reverting to a ‘normal’ phenotype after removal of the antibiotic challenge. Recent efforts to study persisters have involved proteomic and metabolomic analysis[20], imaging the population *in situ*[21], separating persisters using microfluidics [22], and/or its phenotyping with advanced genomic techniques [14, 23, 24] (such as specific gene knockout methods[25], and/or the use of fluorescent reporter proteins), flow cytometry [26], and heavy isotope labelling-mass spectrometry.[27] These techniques are, however, all complicated by the presence of very large numbers of dead/dying cells during the experiments, as well as the cells reverting back to their non-persister form during isolation and analysis. This complication makes it difficult to distinguish the signal of “true” persister genes.

Despite their limitations, these earlier studies have yielded valuable information on the metabolic state during persistence in different species, together with a diverse set of the genes and functions involved, such as type II toxin/antitoxin (TA) module[28, 29], protein degradation systems, purine metabolism, amino acid metabolism and metabolic regulators[30, 31], DNA damage repair[32, 33], SOS response[34, 35], cellular communication and quorum sensing[36, 37], stringent response and ppGpp[38, 39], and efflux pump systems.[40, 41] The lack of clear mechanistic consensus has challenged efforts to understand the causal basis for persistence and to identify targets for their elimination. Our aim in this paper is to develop a novel method to analyze the proteome of persister cells in the Gram-positive pathogen *S. aureus. S. aureus* is is the causative agent for a variety of chronic and relapsing infections, including hard-to-treat sub-dermal and systemic infections.[42-45] It is a clinically problematic species due to the existence of highly drug-resistant strains, such as Methicillin-resistant *Staphylococcus aureus* (MRSA). The widespread distribution of these drug resistant *S. aureus*-strains has led to the death of over 1 million patients a year worldwide[46, 47], with this number predicted to increase rapidly in the coming decades. This species has therefore been designated as a “pathogen of major concern” by the WHO. [43, 48, 49]

Although there have been recent efforts to characterize the persister proteome of *S. aureus*, this work has faced some of the same challenges as other methods, namely the fact that persisters are exceedingly rare. To overcome these limitations, we hypothesized that bioorthogonal, or “click”, chemistry would allow deeper protein coverage of rare persister cells, because it enables retrieval and isolation of the (very rare) proteome during the various stages of persistence followed by characterization of only the proteins that were expressed in persister cells.

Click reactions are a family of high-yielding, fast, and selective chemical ligation reactions that can be performed with a high degree of selectivity in biological environments.[50] They have been used extensively to, for example, image the location of specific lipids in cells [51, 52], look at surface regulation of carbohydrates [53], to quantify DNA-synthesis in cells[54-56], and to study nutrient channel activities.[57, 58] The approach has also been used to label the expressed proteome across a given time window. This approach, called Bio-Orthogonal Non-Canonical Amino acid Tagging (BONCAT)[59, 60], makes use of the fact that in many species, the methionine tRNA/tRNA-synthase pair can accept and incorporate unnatural amino acids with click-reactive chemical groups.[61, 62] The two best mimics of methionine are azidohomoalanine (Aha, **Figure 1a**) and homopropargylglycine (Hpg), which can be both be ligated using copper-catalyzed [3+2]-cycloaddition reactions, which is a very low-background click reaction.[60, 61, 63-65] When cells are pulsed with these amino acids, they are incorporated into the proteome of translationally active cells only during the pulse period, thereby enriching the proteome for the specific set of proteins expressed during a short time window. Here we report using BONCAT-labelling[59, 60][66, 67] to identify expressed proteins in persisters cells that form during the exposure of two different MRSA and VRSA strains to oxacillin or moxifloxacin.

**Figure 1.**
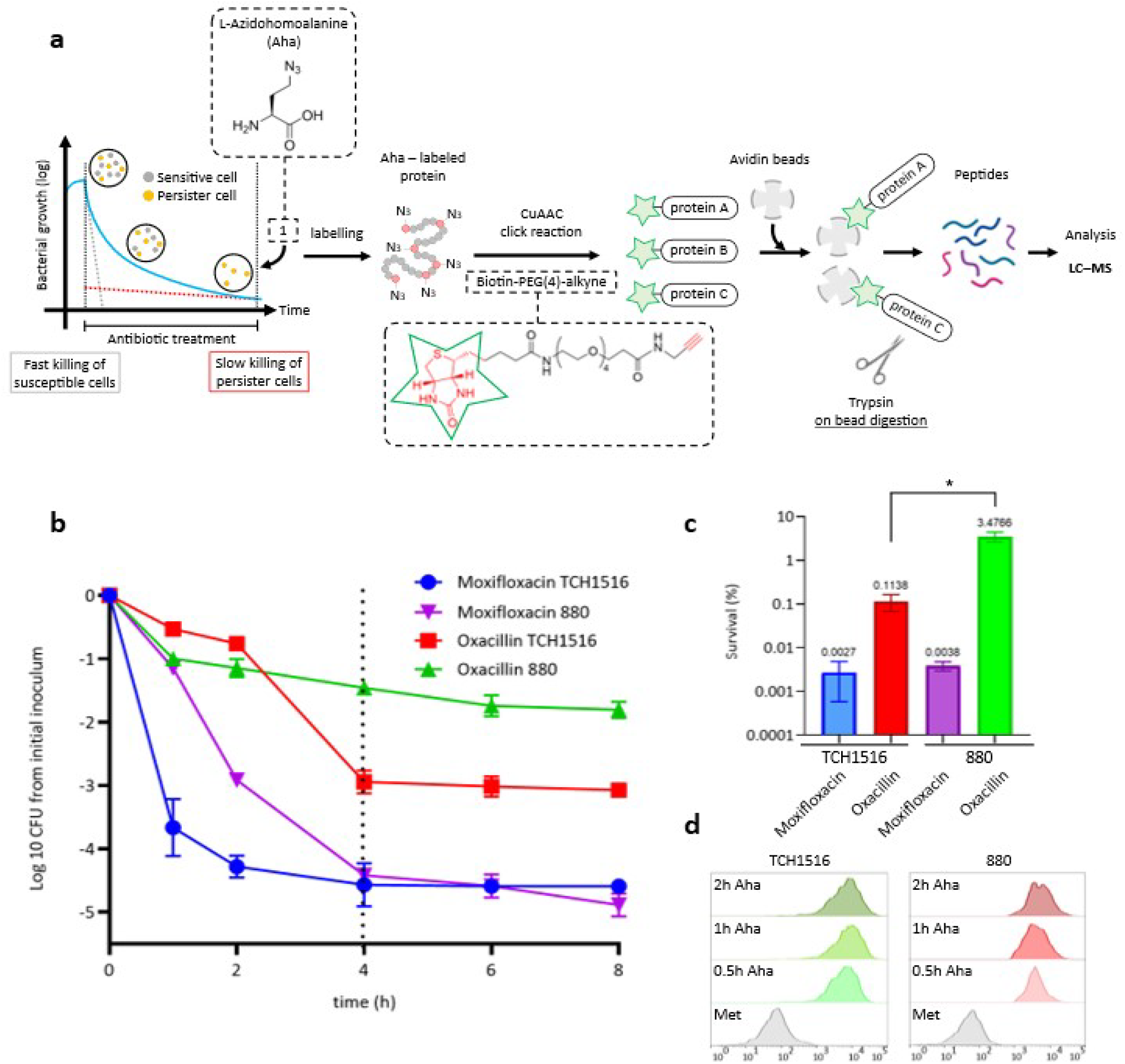
Labelling the expressed proteome of persister populations using BONCAT. **a**. Bioorthogonal labelling of persister cells with click handles to perform click chemistry followd by an avidin-pulldown to selectively acquire labelled proteins for further analysis with LFQ-LCMS based proteomics. **b**. Time Kill Curves of S. aureus strain TCH1516 and 880 during challenge with 50x MIC for Oxacillin and Moxifloxacin (mean <u>+/-</u>SEM, n=3). **c**. Survival (%) for both strains after 4h treatment with 50x MIC Oxacillin and Moxifloxacin (n=4). **d**. Flow cytometric profile of bacteria incubated with Aha. Label incorporation gives positive signal after 30 min incubation.

Characterizing persistence in two different strains with two antibiotics that vary in their mechanism-of-action (one a β-lactam that targets cell wall biosynthesis and the other a fluoroquinolone that targets DNA gyrase) made it possible to identify strain and antibiotic-specific persistence mechanisms. Further, by comparing the proteins expressed during the persister state to the expression profile shortly after antibiotic challenge, we could monitor temporal changes in protein expression as bacteria enter the persister phase.

After confirming the suitability of the BONCAT approach, we found extensive strain and antibiotic-specific changes to the proteome during antibiotic exposure, as well as several shared responses. Among these, we observed that during the induction of the persister state, oxacillin elicited the upregulation of 75 and 33 proteins for TCH1516 (USA300) and 880, respectively, with a downregulation of 129 and 426 proteins, respectively. Moxifloxacin showed the upregulation of 105 and 79 proteins for TCH1516 and 880, respectively, with a downregulation of 151 and 504 proteins respectdively. Within these proteins, we observe the upregulation of the stringent response, cell wall biosynthesis, purine metabolism, ppGpp biosynthesis, two component system (TCS), lipid metabolism, ABC transporters, D-alanine biosynthesis and L-proline degradation. Additionally, we observed the downregulation of proteins associated with amino acid biosynthesis and degradation, protein biosynthesis and modification, carbohydrate metabolism, among others. These results suggest that translation activity of persisters reflects an active defense against antibiotic pressure for survival.

## Results

### Method optimization

To study the persister proteome, we picked two strains of MRSA and two antibiotics: a β-lactam that targets cell wall biosynthesis (oxacillin)[68], and a fluoroquinolone that inhibits DNA-gyrase (moxifloxacin).[69] Strain TCH1516 was chosen for its methicillin resistance[70], and 880 for its broad resistance, including to vancomycin.[71] We first determined the minimum inhibitory concentrations (MIC) of each antibiotic-strain combination. We found the MIC-values for strain TCH1516 to be 10 mg/L for oxacillin and 0.08 mg/L for moxifloxacin. Strain 880 had MIC-values of 20 mg/L for oxacillin and 2.5 mg/L for moxifloxacin. These values are in accordance with those previously reported. [72, 73]

To explore the dynamics of antibiotic-mediated killing we quantified the size of the persister fractions by measuring bacterial cell viability over time using time-kill curves (TKC).[74] Each strain was exposed to 50X the MIC of each antibiotic (for that particular strain), and viability was measured over time (**Figure 1b**).[75, 76] Characteristic biphasic killing for both antibiotics in both strains was observed: an initially rapid drop in CFUs was followed by a second phase around 4h where the decline was far slower.[77, 78] In accordance with earlier TKC-data, we classified the fraction of surviving cells at 4 h as persisters, which varied both for strain and for each antibiotic.[79] The percentages of surviving bacteria were 0.1% for TCH1516 and 3% for 880 upon exposure to oxacillin, and 0.002% and 0.003% for moxifloxacin (**Figure 1c**).

Having established the conditions for inducing persisters of strains TCH1516 and 880 with moxifloxacin and oxacillin, we next determined whether a BONCAT-approach could be developed that could identify whether these strains were translationally active, and whether the technique was sensitive and selective enough so that the translated proteome could be retrieved and identified from this minority fraction of cells.

We first determined whether BONCAT could be used to label the proteome of the two MRSA strains, as BONCAT had only previously been performed on mixtures of bacteria containing *S. aureus*, and not on isolated populations.[80] To do this we determined whether the bioorthogonal amino acid L-azidohomoalanine (Aha) was incorporated in unchallenged 880 MRSA. The cells were incubated with 4 mM of this amino acid, in line with the concentration range used to label other species.[59, 81-83]

To visualize the amount of Aha uptake by the bacteria, the Cu-catalysed Huisgen cyclo-addition click reaction (CCHC) was performed on fixed and permeabilized cells.[84] This reaction leads to the specific ligation of a fluorophore to the azide-residues of Aha, with very little background staining and can thus be used to analyze the uptake of the amino acid on a per cell basis using flow cytometry (**Figure 1d**). These experiments showed an approximately 70-fold increase in fluorescence after 0.5h incubation with Aha.

These experiments did not, however, show whether the amino acid was also incorporated into the proteome as shown for other species.[59, 85-87] To determine whether this was the case, the cells were lysed, subjected to CCHC with an alkyne-Alexa488 fluorophore and analyzed by fluorescent SDS-PAGE. Presence of a fluorescent band at a given MW would indicate the presence of an Aha-labelled protein at that weight. These gels (**Figure S1**) showed incorporation of the azide into proteins over the whole proteome, with the first detectable signal observed after 5 minutes incubation with signal strength increasing over time until 1.5h.

Having confirmed the incorporation of Aha into the nascent MRSA-proteome, we next assessed whether labelling levels were sufficient to also retrieve and analyze the expressed proteome by mass spectrometry, using biotin-PEG-alkyne (**Figure 1a**) to label the azides and a avidin resin to selectively pull down the click-proteins.[88] After incubation of the cells for 1.5h with Aha, followed by lysis and this click pulldown protocol, the retrieved proteins were digested with trypsin and subjected to LFQ-LCMS analysis.[89-92] In total, 1533 proteins were identified. Direct comparison with LFQ-LCMS of trypsinized lysate showed that >90 % of the detected whole proteome was recovered. This confirmed the suitability of the BONCAT-MS approach for retrieval of the expressed MRSA proteome (**Figure 2**), without bias stemming from the Aha-labelling. Label-free quantifications (LFQs) correlate between the Aha and wt MRSA-proteome, suggesting Aha does not skew expression in the species.

**Figure 2.**
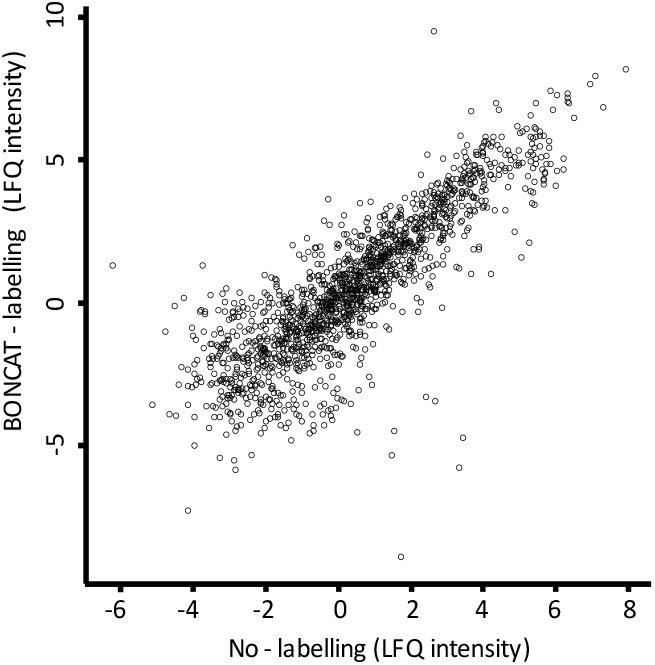
Evaluation of BONCAT-MS on MRSA strain. Correlation between LFQ intensities for BONCAT retrieved proteins compared to the recovery of non labelled proteins. The positive correlation shows that both methods are effective at obtaining MRSA proteome. The comparison shows a linear trend with a Pearson’s correlation score of 0.8.

### BONCAT-MS of persister-expressed proteins

We next applied BONCAT to the isolation and identification of proteins expressed during persistence. Strains TCH1516 and strain 880 were treated with 50x the MIC of oxacillin and moxifloxacin for 4 hours as above, followed by the addition of Aha for 1.5 hours. After this time, the cells were lysed and fluorescent SDS-PAGE was used to check whether detectable levels of Aha-positive proteins were produced during this period. The observed signal confirmed the previously reported translational activity of the persister population (**Figure S2**).[4, 93-96]

### Drug specific effects

In order to determine the changes in protein expression for each strain/antibiotic combination at the 4h timepoint, the treated cells were subjected to the click-pulldown-MS protocol. Multiple different proteins were found to be up-, or downregulated in each of the antibiotic-strain combinations, with very little overlap in the changes in expression between all conditions (1 up- and 22 downregulated) pathway identification of the known proteins identify a common downregulation of purine metabolism (purH, purM, purD, purN, serS)(**Figure 3c**). More similarities were observed within each strain, or upon treatment with the same antibiotic. However, the majority of changes after antibiotic exposure were strain or drug-specific. Of the shared responses, a far greater fraction of proteins were downregulated.

**Figure 3.**
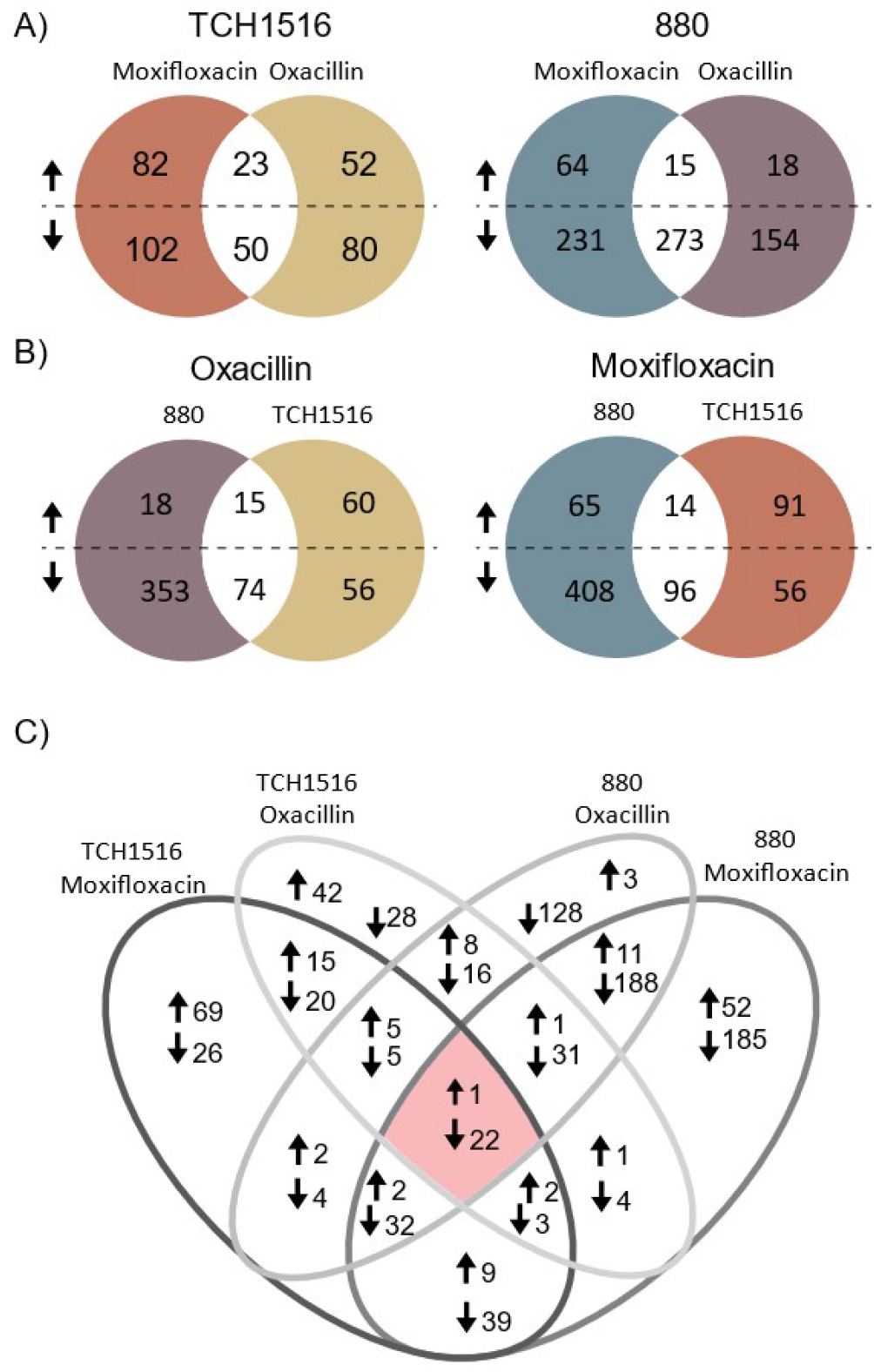
Venn diagram for upregulated and downregulated shared proteins for TCH1516 and 880 after 4h treatment with Oxacillin and Moxifloxacin. **A**. Strain-specific upregulation: TCH1516 has 23 proteins upregulated and 50 downregulated shared between antibiotic challenges. Strain 880 has 15 proteins upregulated and 273 downregulated independent of antibiotic choice. B) Oxacillin induces upregulation of 15 proteins and the downregulation of 74 proteins common to both strains. Moxifloxacin results in the common upregulation of 14 proteins and the downregulation of 96 proteins between both strains. C) Full Venn diagram for upregulated and downregulated shared proteins for TCH1516 and 880 after 4h treatment with Oxacillin and Moxifloxacin.

β-lactam treatment, for example (**Figure 3b, bottom left side**), led to upregulation of 15 shared proteins in both strains. Pathway identification of the known proteins revealed them to be involved in cell wall biosynthesis (mgt, uppS, ptr), TCS (vraS) and *S. aureus* infection (efb). 74 proteins were downregulated in both strains, involved in amino acid degradation (hutG, ald1, tdcB), nucleotide-sugar biosynthesis (glmU), purine metabolism (purH, purM, purD, purN, serS), aminoacyl-tRNA biosynthesis (serS), carbohydrate degradation (Fda, rpiA), fermentation (ldh2), pyrimidine metabolism (pyrE, pyrC) and porphyrin-containing compound metabolism (hemB) (**Table 1, Figure 3b**).

**Table 1.**
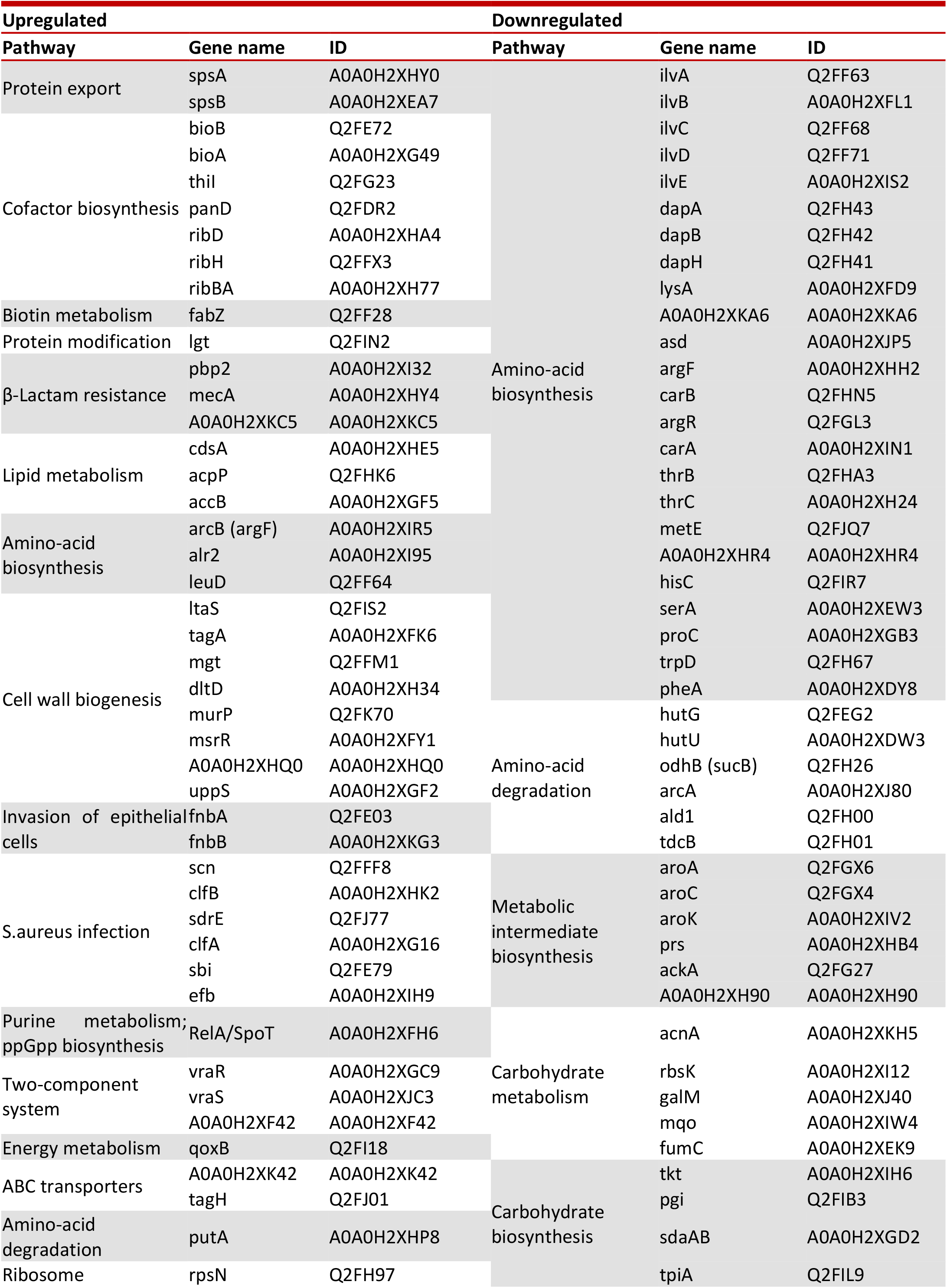
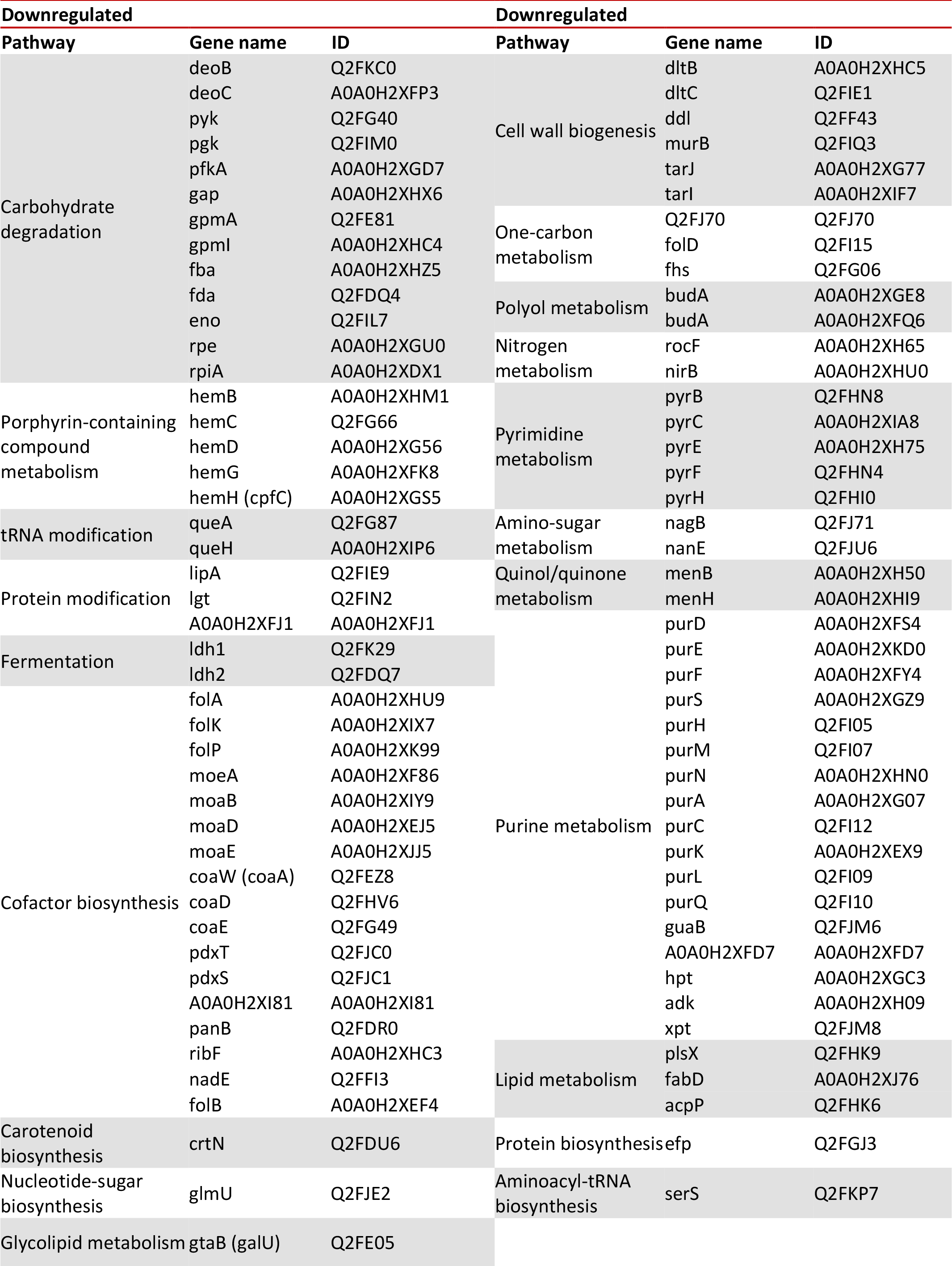
Proteins and pathways upregulated or downregulated in TCH1516 and 880 persisters.

Fluoroquinolone challenge (**Figure 3b, bottom right panel**) led to the upregulation of 14 shared proteins, which were found to be involved in lipid metabolism (accB), and cofactor biosynthesis (ribBA). 96 proteins were downregulated in both strains, involved in purine metabolism (purH, purM, purD, purN purl, purQ, purC, purF, purS), aminoacyl-tRNA biosynthesis (serS), protein biosynthesis (efp), one-carbon metabolism (folD), carbohydrate metabolism (tpiA, acnA), and amino-acid biosynthesis (dapH, dapB, dapA, serA, proC, thrC, ilvE, asd, A0A0H2XHR4) (**Table 1, Figure 3b**).

### Strain specific effects

Some shared changes were found within each strain that suggest common persister pathways for each strain that are not drug dependent. TCH1516 treated with Oxacillin and Moxifloxacin led to the joint upregulation of 23 proteins of which the known proteins could be allocated to the TCS (vraR, vraS, CytD), purine metabolism and ppGpp biosynthesis (RelA/SpoT), lipid metabolism (acpP), *S. aureus*-infection (efb) and cell wall biosynthesis (ptr). 50 proteins were downregulated in common, involved in pathways of amino-acid biosynthesis (ilvA, ilvB, A0A0H2XKA6), purine metabolism (purH, purM, purD, purN) and aminoacyl-tRNA biosynthesis (serS) (**Table 1, Figure 3a**).

Strain 880 showed the common upregulation of 15 proteins related to the ABC transporter family (A0A0H2XK42, tagH), cell wall biosynthesis (mecA), D-alanine biosynthesis (alr2), *S. aureus* infection (sbi) and L-proline degradation (putA). 273 proteins were downregulated upon treatment with both antibiotics, of which the known ones could be allocated to carbohydrate degradation pathways (Fda, rpiA), fermentation (ldh2), amino acid degradation (ald1, tdcB), pyrimidine metabolism (pyrE, pyrC), porphyrin-containing compound metabolism (hemB), purine metabolism (purH, purM, purD, purN, purl, purQ, purC, purF, purS), aminoacyl-tRNA biosynthesis (serS), protein biosynthesis (efp), one-carbon metabolism (folD), carbohydrate biosynthesis (tpiA), carbohydrate metabolism (acnA) (**Table 1, Figure 3a**).

Most proteins identified have not yet been functionally annotated. A total of 1502 proteins were measured for TCH1516 and 880 under Oxacillin and Moxifloxacin treatment, of which 20% hits were successfully found in reported pathways, including 4% upregulated and 16% downregulated proteins. The function of the remaining 80% of proteins remains unknown.

### Temporal changes in the TCH1516 proteome under moxifloxacin treatment

Having identified the proteomic profile of the TCH1516 and 880 persister cells after 4h of antibiotic exposure, we next wanted to study how the expressed proteome changed over time as cells went from the early “stressed” stage to the slow-dying “persister”-like state. For this, we chose to treat the highest persister-fraction strain (TCH1516) with moxifloxacin and apply the Aha-pulse after 5 min, 0.5h, 1h, 1.5h, 2h, 3h or 4h after the initial antibiotic challenge. The pulse length of 1.5h was maintained. At the end of the Aha pulse, the bacteria were lysed, and either subjected to a CCHC reaction with a fluorophore for SDS-PAGE visualization or reacted with biotin alkyne for protein enrichment and analysis with LCMS. Fluorescent SDS-PAGE analysis of the lysed bacteria (**Figure S3**) showed a decrease in the total amount of protein labeled over time, as well as a different protein expression fingerprint over time, suggesting proteome remodeling promoted by the bacteria while entering a persistent state.

A comparison between the fold change of each of the significantly changed proteins compared to the untreated controls was performed (**Supplementary table 1**). The heat map derived from this comparison shows a broad decrease in protein expression in the first 1,5 hours after starting treatment, with a gradual increase in expression observed as the bacteria move towards the persister state (**Figure 4a**). To focus on the clearest changes through time, the expression profiles at 5 min were compared to those at 4h after antibiotic challenge (**Figure 4b**). This difference was particularly striking, with the expression of 211 up and 30 downregulated proteins. Of these, 200 and 27 respectively are of as yet unknown function (**Supplementary table 1**). Proteomic results from this comparison from known pathways or with highly significant up or down regulation are shown through time in **Figure 4c**. As with the 4h timepoint, we observed the downregulation of the targets mapped to purine metabolism (purC, purS), as well as upregulation of proteins mapped to lipid metabolism (acpP, accB), amongst others.

**Figure 4.**
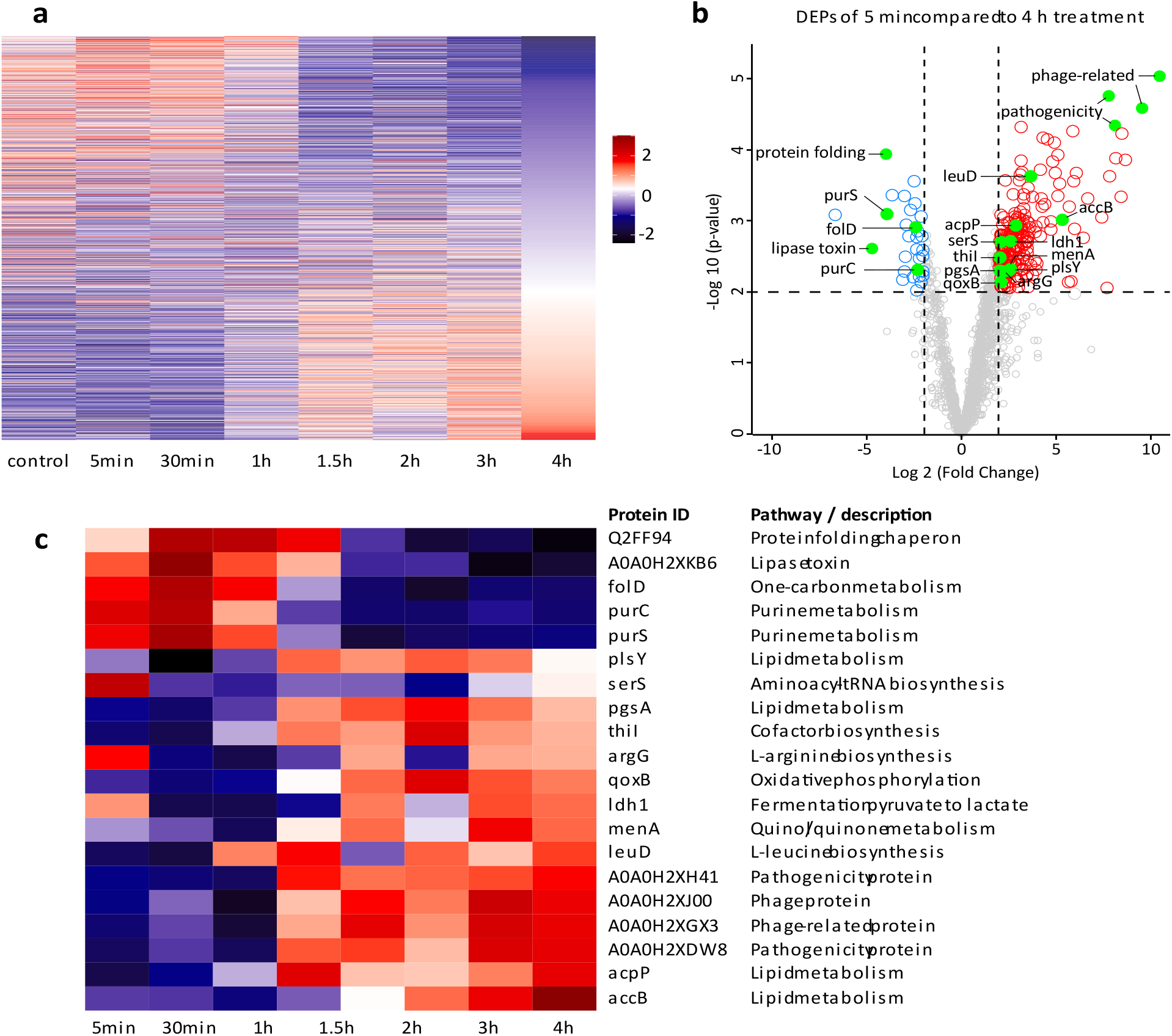
Protein diversity expression of S. aureus TCH1516 during moxifloxacin treatment. a. Heatmap displaying all the proteins identified across antibiotic treatment (n=4). b. Differentially Expressed Proteins (DEPs) proteins after 4h compared with initial phase of the treatment at 5 min. Comparison revealed 211 upregulated and 30 downregulated proteins from which 200 and 27 are unknown function. Additional proteins were selected due to its high significance such as A0A0H2XJ00 and A0A0H2XGX3, were found to be phage proteins, A0A0H2XH41 and A0A0H2XDW8 were identified as pathogenicity proteins, A0A0H2XKB6 is a lipase toxin and Q2FF94 a chaperon folding protein. c. Heatmap displaying the DEPs identified in b for proteins with known pathways and with high significance for S.aureus TCH1516 at different time points with moxifloxacin treatment. Pathway identified proteins are listed in supplementary table 1.

## Discussion

To our knowledge, this study is the first example to use BONCAT to retrieve and identify the persister proteome. Two strains of *S. aureus*, TCH1516 and 880, were challenged with a cell wall disrupting antibiotic (oxacillin) or a gyrase inhibitor (moxifloxacin). The reason was to identify shared or unique mechanisms that regulate the persister phenotype. First, we confirmed that persisters are translationally active. Next, consistent with other observations, we found that up- and down-regulated proteins in persisters are highly strain and antibiotic specific. [47] This suggests that single diagnostic targets for persister cells may remain elusive. Our results also suggest the value of a BONCAT approach in identifying proteins that may be causally involved in regulating this phenotype. [18, 97] The depth and breadth of coverage during persistence allowed us to identify changes in expression of hundreds of known and unknown proteins. These data therefore provide an exciting starting point to begin to elucidate of the mechanism(s) by which bacteria can enter the persister state. Importantly, BONCAT that only antibiotic stress, and not “BONCAT-stress” was howed no interference on bacterial growth, ensuring responsible for the observed changes.

The mapping of the known proteins to universal databases such as Uniprot or KEGG highlights current limitations in understanding persistence. The retrieval of 80% unknown proteins means that we are likely only beginning to scratch the surface of this complex phenomenon. Nevertheless, some common themes emerge from our assays and some of the known pathways overlap with other studies. Both strains under different antibiotic challenges revealed the essential role of maintaining cell wall integrity, with the joint expression of TCS to support cell-wall damage related pathways.[98, 99] This finding is consistent with results from other groups, where In response to the cell wall targeting antibiotic *S. aureus* persisters increased cell wall biosynthesis related proteins, such as those involved in the production of wall techoic acids (ltaS, tagA, dltD, msrR, A0A0H2XHQ0) and peptidoglycan (mgt, murP, uppS).[100] Upregulation of ABC transporters, as observed in this dataset, has also been observed by others and is believed to indicate increased efflux to remove antibiotics from exposed cells.[53, 101]

We observed changes in several pathways associated with regulating cellular metabolism and replication. This result is supported by the decrease in proteins in anabolic pathways also observed by Liu *et al*., 2024.[20, 100] The activation of stringent response driven by ppGpp in response to nutrient limitation (*e*.*g*., carbon, amino acids, nitrogen, phosphate) was also seen, in alignment with previous reports, where similar characteristics were reported for *S. aureus* persisters in response to antibiotic damage.[20]

This response has been reported to trigger bacterial dormancy through downstream signaling of pathways involving toxin-antitoxin (TA) modules[102, 103], leading to a general shut down of vital activities in response to stress.[104] These observations, and their alignment with the previous reports of persister biology lend further credence to the BONCAT-approach.[105, 106] Bacterial strain and antibiotic class were found to have a profound influence on persister formation. Upregulation compared to the unchallenged control of proteins involved in the teichoic acid pathway, cell wall damage (pbp2) and activation of beta-lactam resistance (mecA) specifically in the MRSA strains stressed with oxacillin, shows the antibiotic-specific phenotypic traits displayed by surviving population. [107-109] Thus, resistance and persistence as observed in the context of the study, seem to be two different and complementary adaptations exhibited by bacteria in response to antibiotic exposure. They can co-exist and play synergistic roles in bacterial survival under antibiotic pressure. Antibiotic strategies should therefore take both resistance and persistence into account.[110, 111]

A major surprise during this study was the large number of as-yet unidentified highly enriched proteins. Indeed, the majority of changed proteins during the moxifloxacin time course are uncharacterized. Although these point to multiple paths for future experiments, they also highlight that it may be difficult to identify common pathways to target persister cells arising in different strains, even of the same species.[112]

In all, this deepened proteomic recovery of the persister population can, in combination with application to other species, hopefully aid to identify new potential targets for treating this most troublesome of infection phenomena.

## Materials and methods

### Bacterial strains and growth conditions

The bacterial culture strains were US300 TCH1516 and 880; BR-VRSA. Mid-exponential phase cultures were prepared by diluting overnight cultures with 1:50 in Luria−Bertani (LB) broth and incubating these at 37 °C at 100 rpm until the optical density (OD600 nm) had reached 0.4.

### Chemicals

All antibiotics were purchased from Sigma-Aldrich and used without further purification. Aha was purchased from Click Chemistry Tools. Neutravidin beads (Thermo Scientific Pierce, catalog number 20219)

### Minimum Inhibitory Concentration

MIC determination was done on 96 well plate setup in biological triplicates. Mid-exponential phase cultures were prepared as above. Prior to the experiment, they were diluted to a starting inoculum size of 10^6^ CFU/mL. They were transferred to a 96-well plate containing Mueller-Hinton broth and antibiotic solution at a desired concentrations to yield a final inoculum concentration of 5 × 10^5^ CFU/mL and incubated for the indicated time. The content of the wells was plated out on LB-Agar plates, which were incubated overnight at 37°C. The MIC was determined as the lowest concentration at which no visible bacterial growth was observed.[72, 73]

### Time-Kill assay

Mid-exponential phase cultures were prepared as above and were diluted to a starting inoculum of 10^6^ CFU/mL. They were then exposed to the antibiotics at a final concentration of 50x the MIC[78], and incubated at 37°C, 100 rpm. Samples at different times were taken, where necessary diluted in PBS (when too many colonies had formed for counting), performed, then plated on LB agar plates and CFUs were counted after overnight incubation at 37°C.[113]

### BONCAT labelling and lysis

Sample preparation was performed according to the procedure described in [64, 82]. BONCAT labelling of the persisters was done the following way: bacteria were exposed to antibiotic in LB-medium for the indicated time, after this, the bacteria were pelleted by centrifugation for 5 minutes at 3000 *rcf*. The pellet was resuspended in SelenoMet medium (from Molecular Dimensions) augmented with the antibiotic at the same concentration in order to ensure that the bacteria remains under stress at all times. The samples were left on ice for 5’ following which they were centrifuged at 14000g at 4°C for 10, prior to resuspension in fresh SelenoMet medium augmented with 4 mM Aha.[114] The cells were incubated for 1.5 h, after which the cells were harvested by centrifugation and resuspended in in lysis buffer (PBS, 4% SDS, with Roche EDTA-Free Protease Inhibitor added as per manufacturer’s recommendation). The cells were lysed using a Bead Homogenizer (MP FastPrep-24 5G). The a Bead Homogenizer was ran for 10 cycles of 50 seconds each at 6 m/s, with 3-minute intervals on ice in between cycles. Following homogenization, tubes were centrifuged at 1500 *rcf* for 2 minutes and supernatant was transferred to clean tubes. These were centrifuged again for 30 minutes at 4°C at 14000 rpm, sterile-filtered through 0.2 μm membranes. Next the reduction-alkylation was performed by incubation with DTT (1M) was added per 1 mL sample, incubated for 15 minutes at 65°C, 600 rpm. Next 80 μL of IAA stock (0.5M) was added and incubated for 30 minutes at room temperature in darkness, next 350 μL of SDS stock (10%) was added and incubated for 5 minutes at 65°C. Next samples followed Methanol/Chloroform precipitation.[115] BCA assay was performed according to the manufacturer’s protocol, to determine the protein concentration.

### Analysis of BONCAT Labelling by flow cytometry

*S. aureus* was metabolically labelled as described above and samples with OD600 ≈ 0.4 were collected after 30 minutes, 1h and 2h to analyze the label incorporation levels at single bacterium level by flow cytometry. Bacterial samples were pelleted by centrifugation (10 min at 5000 *rcf*), washed once with PBS and resuspended in 100 μL 4% PFA for overnight fixation.

Fixed bacteria was washed once with 100 µL cold PBS and centrifuged 10 minutes at 6000 *rcf* then permeabilized in 50 µL permeabilization buffer (0.1% Triton-X100 in PBS) for 20 minutes. Permeabilized bacteria was washed once with 100 µL PBS and incubated with 50 µL of 1x concentrated click mix for 1h in the dark at RT (table 1). After incubation, cells were washed once with 100 µL PBS and resuspended in 100 µL Washing Buffer (1% BSA in PBS) and incubated for 30 minutes in the dark at RT. After incubation, cells were washed once with 100 µL FACS buffer (EDTA 2mM in PBS) and resuspended in 200 µL FACS buffer, then flow cytometry analysis was performed. Aha-AF488 was detected in the FITC channel. The analysis was performed using the guava InCyte software and all subsequent analysis were performed with FlowJo V10.7.2 (FlowJo software). The measured events were gated on size, shape and fluorescence to accurately select single bacteria.

**Table 1.**

| Table 1 |  |
| --- | --- |
| Chemical | Final Concentration |
| CuSO<sub>4</sub> | 1 mM |
| Sodium ascorbate | 10 mM |
| THPTA ligand | 1 mM |
| Aminoguanidine | 10 mM |
| Hepes pH 7.4 | 96 mM |
| AF488-alkyne | 5 μM |

### Fluorescence SDS-PAGE analysis

20 μg of protein was diluted to a final volume of 10 μL with HEPES-buffer (100 mM pH 7.4). 5 µL of a 3x concentrated mixture of click reagents (table 2) was added, and the mixture incubated for 1h at RT in the dark. Sample buffer without thiol was added, the sample heated to 95 °C for 10 minutes, followed by SDS-PAGE separation. Prior to Coomassie staining, the gels were imaged in a ChemiDoc fluorescent gel scanner with the 700/50, 602/50 or 532/28 nm filter set.

**Table 2.**

| Table 2 |  |
| --- | --- |
| Chemical | Final Concentration |
| CuSO<sub>4</sub> | 3 mM |
| Sodium ascorbate | 30 mM |
| THPTA ligand | 3 mM |
| Aminoguanidine | 30 mM |
| Hepes pH 7.4 | 300 mM |
| AF488-alkyne | 15 μM |

### Bioorthogonal pull-down

Sample preparation was performed according to the procedure described in [92]. 300 μg of protein was volume adjusted to 2 mg/mL (150 µL). An equal volume of 2x concentrate of click mix was added (table 3), and the mixture was reacted 2h at RT in the dark under gentle rotation.

**Table 3.**

| Table 3 |  |
| --- | --- |
| Chemical | Final Concentration |
| CuSO<sub>4</sub> | 2 mM |
| Sodium ascorbate | 20 mM |
| THPTA ligand | 2 mM |
| Aminoguanidine | 20 mM |
| Hepes pH 7.4 | 200 mM |
| Biotin-PEG-alkyne | 160 μM |

Excess unreacted biotin-PEG-alkyne tag was removed by precipitating out the proteins with chloroform/methanol. First 200 µL of 50 mM Hepes (100 mM, pH 7.4) was added, followed by 666 µL methanol. After vigorous vortexing, 166 µl chloroform was added, followed by a further burst of vortexing. 150 µL of water was added to the mixture to cause phase separation. Centrifugation for 10 min at 10,000 *rcf* at room temperature yielded a three layer system where the top layer was water/methanol, the white film at the interface is the precipitated protein and the bottom phase the chloroform/methanol. The top layer was removed carefully, 600 µl methanol was added and mixed gently followed by centrifugation 10 min at 10,000 *rcf* and RT. After this, the supernatant was removed and the pellet dried for <2 min in air. The pellet was resuspended in 250 µl urea buffer (8 M urea in 25 mM ammonium bicarbonate, pH 8.0).

Next, 100 μL of Neutravidin beads (Thermo Scientific Pierce, catalog number 20219) were washed three times with 250 μL PBS and resuspended in 2 mL PBS, the proteins were added on top and incubated for 3 hours at room temperature under gentle rotation. After which the beads were collected by centrifugation (2 minutes at 2500 *rcf*) and washed 5 times with 2 mL PBS with vigorous shaking followed by centrifugation to remove any SDS. After the final wash, the supernatant was removed and 250 μL of on-bead digestion buffer (100 mM Tris pH 8.0, 100 mM NaCl, 1 mM CaCl2, 2% ACN) was added to the each of the bead residues, the beads were transferred to low binding tubes (1.5 mL, Sarstedt). Each sample was treated with 1 μL of trypsin solution (0.5 μg/μL Sequencing Grade Modified Trypsin, Porcine (Promega) in 0.1 mM HCl), and the samples were incubated at 37°C overnight while shaking (950 rpm).

To each sample formic acid (3 μL) was added, followed by filtering off the beads over biospin columns (Bio-Rad, 7326207) on top of 2 mL Eppendorf tubes using centrifugation (2 min, 300 *rcf*). Note: now the 2 mL Eppendorf tubes contain the main sample solution. Next, StageTips were used for subsequent desalting of the samples according to the procedure described in [116]. StageTips were placed in holders in Eppendorf tubes to collect the flow after each individual step which is followed by centrifugation (2 min, 300 *rcf*). The conditioning of the StageTips started with 50 μL MeOH, washing with 50 μL solution B (80% acetonitrile v/v, 0.5% v/v formic acid in water), and 50 μL solution A (0.5% v/v formic acid in water). The sample solution was loaded through the StageTips, followed by a wash with 50 μL solution A. The StageTips were then transferred to low binding tubes, and the tips were flushed with 100 μL solution B. The collected sample was concentrated in a SpeedVac (45°C, V-AQ, 1-2 hrs) (Eppendorf Concentrator 5301). The samples were stored at 20°C until LC-MS measurement.

### LC-MS measurement and analysis

Samples were reconstituted in 50 μL LC-MS sample solution (3% v/v acetonitrile, 0.1% v/v formic acid in Milli-Q) to follow Nanodrop measurement. Samples were diluted to 100 ng/μL in LC-MS sample solution containing 10 fmol/μL yeast enolase digest (cat. 186002325, Waters). Injection amount was titrated using a pooled quality control sample to prevent overloading the nanoLC system and the automatic gain control (AGC) of the QExactive mass spectrometer. The desalted peptides were separated on a UltiMate 3000 RSLCnano system set in a trap-elute configuration with a nanoEase M/Z Symmetry C18 100 Å, 5 µm, 180 µm x 20 mm trap column (Waters) for peptide loading/retention and nanoEase M/Z HSS C18 T3 100 Å, 1.8 µm, 75 µm x 250 mm analytical column (Waters) for peptide separation both kept at 40 °C in a column oven.

Samples were injected on the trap column at a flow rate of 15 µL/min for 2 min with 99% mobile phase A (0.1% FA in ULC-MS grade water (Biosolve)), 1% mobile phase B (0.1% FA in ULC-MS grade acetonitrile (Biosolve)) eluent. The 85 min LC method, using mobile phase A and mobile phase B controlled by a flow sensor at 0.3 µL/min with average pressure of 400-500 bar (5500-7000 psi), was programmed as gradient with linear increment to 1% B from 0 to 2 min, 5% B at 5 min, 22% B at 55 min, 40% B at 64 min, 90% B at 65 to 74 min and 1% B at 75 to 85 min. The eluent was introduced by electro-spray ionization (ESI) via the nanoESI source (Thermo) using stainless steel Nano-bore emitters (40 mm, OD 1/32”, ES542, Thermo Scientific).

The QExactive HF was operated in positive mode with data dependent acquisition (no lock mass), default charge of 2+ and external calibration with LTQ Velos ESI positive ion calibration solution (88323, Pierce, Thermo) every 5 days to > 2 ppm. The tune file for the survey scan was set to scan range of 350 – 1400 m/z, 120,000 resolution (m/z 200), 1 microscan, automatic gain control (AGC) of 3e6, max injection time of 100 ms, no sheath, aux or sweep gas, spray voltage ranging from 1.7 to 3.0 kV, capillary temp of 250 °C and an S-lens value of 80 V. For the 10 data dependent MS/MS events the loop count was set to 10 and the general settings were resolution to 15,000, AGC target 1e5, max IT time 50 ms, isolation window of 1.8 m/z, fixed first mass of 120 m/z and normalized collision energy (NCE) of 28 eV. For individual peaks the data dependent settings were 1.00e3 for the minimum AGC target yielding an intensity threshold of 2.0e4 that needs to be reached prior of triggering an MS/MS event. No apex trigger was used, unassigned, +1 and charges >+8 were excluded with peptide match mode preferred, isotope exclusion on and dynamic exclusion of 10 sec. In between experiments, routine wash and control runs were done by injecting 5 µl LC-MS solution containing 5 µL of 10 fmol/µL BSA or enolase digest and 1 µL of 10 fmol/µL angiotensin III (Fluka, Thermo)/oxytocin (Merck) to check the performance of the platform on each component (nano-LC, the mass spectrometer (mass calibration/quality of ion selection and fragmentation) and the search engine).

### MS data analysis

Raw files from LC-MS measurement were analysed using the MaxQuant software (version 1.6.17.0) with Andromeda search engine.[117] The settings applied for the analysis were as follows: fixed modification: carbamidomethylation (cysteine); variable modification: oxidation (methionine), acetylation (N-terminus); proteolytic enzyme: trypsin/P; missed cleavages: 2; main search tolerance: 4.5 ppm; false discovery rates: 0.01. The options “LFQ” and “match between runs” were checked, while “second peptides” was unchecked. Searches were performed against the UniProt database FASTA file for the *S. aureus* USA300 proteome (Uniprot ID: UP000000793, downloaded 05-03-2023). The data was extracted from “peptides.txt” and “proteingroups.txt” files to obtain protein coverage and MaxQuant scores and for Perseus analysis.

The first analysis of the MaxQuant output was performed using Perseus (version 1.6.15.0). The protein group txt file was loaded on Max Quant, then LFQ intensity entries were selected to main section. The data matrix loaded was filtered by applying filter rows followed by filter Rows based on categorical column and only identified by site option, the resulting matrix was filtered again by applying filter rows followed by filter Rows based on categorical column and reverse option, the resulting matrix was filtered again by applying filter rows followed by filter Rows based on categorical column and potential contaminant option. Next, the categorization of the different files in groups according to experimental design was performed by annotation rows and categorical annotation rows option, in this section all the samples for the same condition were labelled under the same name. Resulting data was transformed into log2 values by choosing basic and transform options. The matrix was filtered by filter rows followed by filter rows based on valid values. To solve the problem of missing values, data imputation was performed with a replacement from normal distribution with width: 0.3 and down shift: 1.8. Normalization was performed by subtraction and change from rows to columns with the subtraction of the most frequent value. With the resultant matrix, volcano plots were performed with the first group as the problem sample and the second group as the control sample. The -log p value and difference cut off was set at ± 2.

The hits above the threshold were obtained from the matrix derived from the volcano plot, this data was used for comparison against UniProt protein database for protein annotation and pathway allocation. The selection of proteins with associated pathway was done using excel. Venn diagrams and were generated using R studio package (7-64), loading the data tables with hits above threshold and the reported pathways previously obtained after uniprot comparison and excel processing. The database for the heatmaps were the result of the additional analysis on Perseus from the database derived from the volcano plots, applying a multiple sample test ANOVA, the matrix was filtered based on categorical column according to ANOVA significant values with the mode selection on keeping matching rows, then a normalization based on Z-score based on rows then a second normalization based on Z-score based on columns, the matrix derived was loaded in excel to average the technical replicates and biological replicates for the same condition and then loaded in R to obtain the heatmaps.

